# Orthosteric interactions with PIP_2_ activate TMEM16A channels

**DOI:** 10.1101/2025.02.24.639951

**Authors:** J Xu, A Santa-Cruz, Aishwarya Chandrashekar, Takeharu Kawano, Mehreen Zaka, Meng Cui, Diomedes E. Logothetis, Leigh D. Plant

**Affiliations:** Department of Pharmaceutical Sciences and Center for Drug Discovery, School of Pharmacy and Pharmaceutical Sciences, Bouve College of Health Science, Northeastern University, Boston MA 02115; Department of Bioengineering, Roux Institute, Northeastern University, Portland ME 04101

**Author notes:** These authors contributed equally to this work (J Xu, A Santa-Cruz).

## Abstract

TMEM16A channels pass Ca^2+^-activated Cl^-^ currents that drive a plethora of fundamental physiological processes. TMEM16A channels are activated by a rise in intracellular Ca^2+^ levels but also require interactions with the signaling phospholipid phosphatidylinositol 4,5-bisphosphate (PIP_2_) to gate open. Although PIP_2_ is essential for the activity of many types of ion channels, its precise binding site and role in channel gating remain poorly understood in most cases, limiting efforts to study channel dynamics and design targeted modulators. In this study, we identify the PIP_2_ binding interactions that govern TMEM16A gating and permeation. Using a combination of gating molecular dynamics (GMD) simulations and electrophysiological assays, we reveal how the α4 helix of TMEM16A interacts with both the phosphate headgroups and acyl chains of PIP_2_ to open an electrostatic ring within the channel and stabilize the extracellular opening of the Cl^-^ conduction pathway. These findings provide key insights into the dynamic role of PIP_2_ in membrane protein function and shed light on the activation mechanism of TMEM16A. This work establishes a framework for rational targeting of TMEM16A in drug development, with potential therapeutic applications in a variety of diseases.

## Introduction

TMEM16A, also known as ANO1, is the prototypical member of the anoctamin family of transmembrane proteins and functions as a Ca^2+^-activated chloride channel ^1–3^. In humans, TMEM16A regulates the electrical excitability of muscle and multiple types of neurons. The channel is also expressed in epithelial tissues where it plays a key role in regulating cell volume, ion homeostasis, and solute transport ^4^. Increasingly, quantitative expression profiling in pathophysiological studies is establishing a link between changes in anoctamin activity, and a variety of pathophysiological processes, highlighting TMEM16A channels as promising druggable targets for several diseases, from hypertension, moyamoya disease to various types of cancer^5,6^.

TMEM16A channels form as dimers of subunits that each contain ten transmembrane helices (α1-10) and a putative Cl^-^ permeation pathway. Under physiological conditions, the channels activate when the intracellular Ca^2+^ concentration exceeds approximately 150 nM, with Ca^2+^ binding at two well characterized sites ^4,7–9^. Despite these foundational insights from Cryo-EM structures, the mechanistic basis for channel gating and the permeation pathway are not resolved.

Several functional studies have demonstrated that, in common with many types of membrane proteins, and most types of ion channels, TMEM16A gating is dependent on the presence of PIP_2_ in the plasma membrane ^10–12^. Although PIP_2_ constitutes only a minor fraction of the plasma membrane, it is increasingly recognized as a master regulator of ion channel function. Despite this pivotal role in membrane physiology, the precise mechanisms by which PIP_2_ regulates channel gating, including that of TMEM16A, are often unclear ^13–15^. *In silico* docking, and mutagenesis studies have identified several potential PIP_2_ binding pockets on TMEM16A; however, the functional significance of these sites is ambiguous ^16^. Furthermore, it is unclear whether these binding sites serve orthosteric roles, directly mediating channel activation, or act allosterically to modulate gating indirectly. This knowledge gap limits our understanding of the channel’s activation mechanics and poses a significant challenge to the rational design of drugs targeting TMEM16A, hindering therapeutic advancements aimed at modulating its activity. Here, we identify the PIP_2_ binding interactions that directly govern how TMEM16A channels are activated. We characterize how PIP_2_ interacts with the α4 and α6 helices of TMEM16A through both its phosphate headgroups and acyl chains revealing the mechanism that underpins channel gating. When Ca^2+^ binds to TMEM16A, PIP_2_ reorients in its binding pocket to facilitate the opening of an electrostatic gating ring within the channel core and stabilize the opening of the extracellular mouth of the ion conduction pathway.

### Identification of the PIP_2_ interface on TMEM16A

Using induced fit docking we identified four distinct PIP_2_ binding pockets (sites-1 to 4) in the model of mouse TMEM16A constructed from the published structure (PDB: 5OYB) (**Fig 1a-b; Fig S1**). Sites 2-4 overlap with PIP_2_ interaction interfaces that have been proposed by others ^10,11,16,17^. Simulating each system using Gating Molecular Dynamics (GMD) revealed that Cl^-^ ions permeated the channel only when PIP_2_ interacted with the site-1 binding pocket. Thus, during 2000 ns simulations, the site-1 system passed 100 ± 48 Cl^-^ per µs (mean ± SD, n = 3) when Ca^2+^ and PIP_2_ were both bound to the channel. No permeation events were observed in the apo-channel or the PIP_2_ alone systems, while relatively fewer permeation events were seen with Ca^2+^ alone (**Fig 1b**). The Cl^-^ ions permeated the channel through a dedicated pathway formed by a series of basic residues including the α4 (R535, R562, K566) and α6 (K645, K661) helices of each subunit (**Fig. 1a**). Consistent with the requirement for both Ca^2+^ and PIP_2_ to gate the channel, principal component analysis (PCA) revealed large conformational changes in the extracellular domain of the α4 helix when both regulators were present. The notable fluctuation in the movement of α4 contrasts with the limited motion of helices α1-7, in the presence and absence of Ca^2+^ and PIP_2_ (**Fig. 1c-e**). A significant decrease in the flexibility of the α6 helix was also observed in the presence of Ca^2+^ (**Fig. 1eii**).

**Figure 1:**
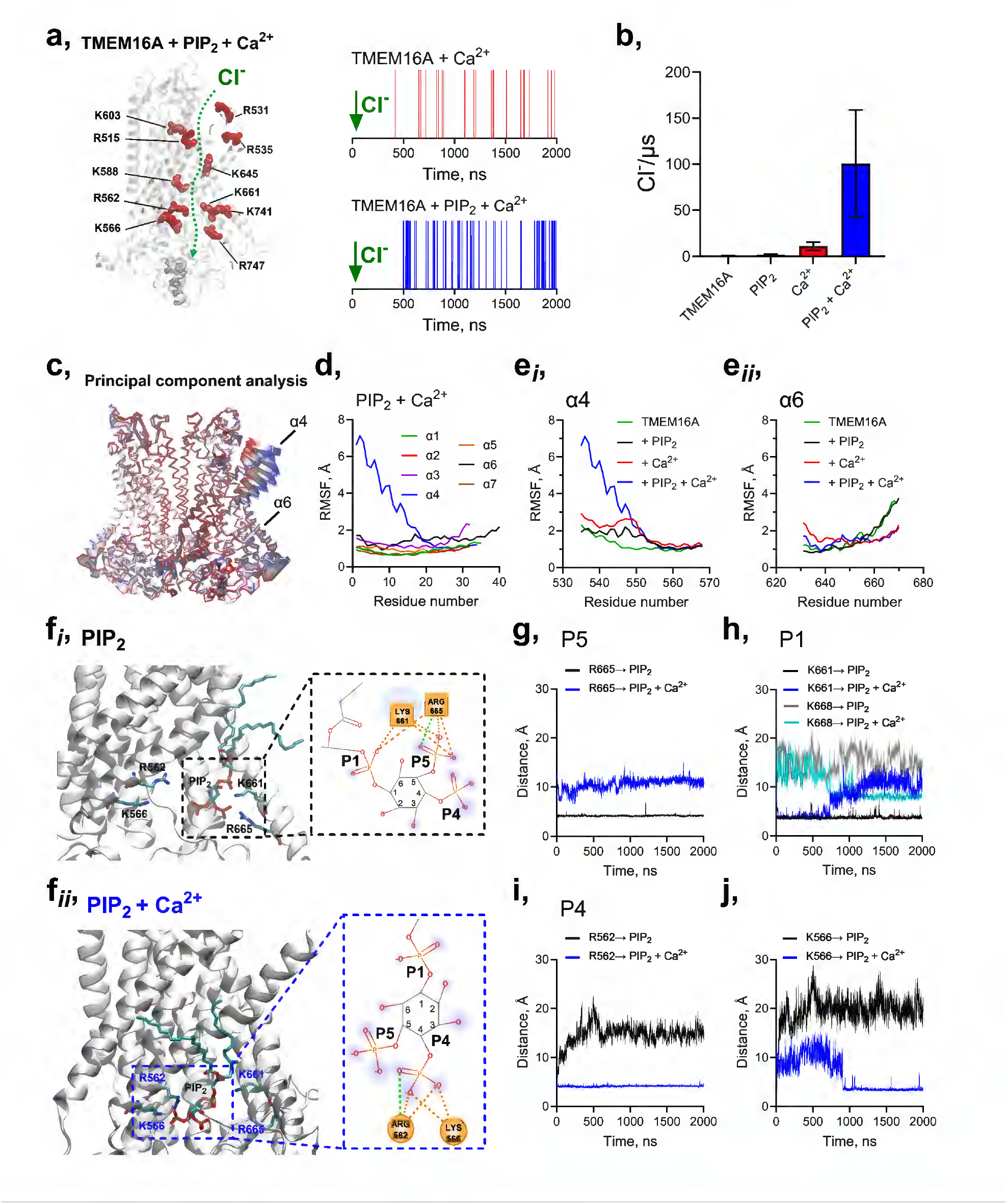
An orthosteric binding site on TMEM16A for PIP_2_. Ca^2+^-bound TMEM16A channels (PDB: 5OYB) activate and pass Cl^-^ *in silico* when PIP_2_ is docked to site-1. **a**, *Left*: The Cl^-^ permeation pathway through a TMEM16A subunit; PIP_2_ and Ca^2+^ are removed for clarity. *Right*: Example raster plots showing Cl^-^ permeation events. **b**, Mean ± S.D. for three simulation replicates measuring Cl^-^ flux per *µs* in different TMEM16A systems (apo, +PIP_2_, +Ca^2+^, and +PIP_2_+Ca^2+^). When present, PIP_2_ is in site-1. **c**, Principle component analysis (PCA) showing increased motion of the external α4 helix in the PIP_2_ + Ca^2+^ system and decreased motion of the internal α6 helix in Ca^2+^-containing systems. **d**, Summary of RMSFs across the length of helices α1-7. **ei**, The fluctuation in α4 motion requires Ca^2+^ and PIP_2_. **eii**, Movement of α6 is subdued in the Ca^2+^-containing systems. **fi**, Binding pose of PIP_2_ in site-1 in the absence and, **fii**, presence of Ca^2+^. **g**, Trajectories of interaction between the P5-phosphate of PIP_2_ and R665. **h**, Trajectories of interaction between the P1-phosphate of PIP_2_ and K661 and K668. **i**, Trajectory of interactions between the P4-phosphate of PIP_2_ and R562 and, **j** K566. *Related to Supplemental Figure S1.*

Under basal physiological conditions, PIP_2_ is available in the plasma membrane to interact with TMEM16A. However, the channel exhibits only low activity until Ca^2+^ ions bind. To investigate the role of PIP_2_ in the gating process, we analyzed the trajectories of interactions between the channel and PIP_2_ in the presence and absence of bound Ca^2+^. In the absence of Ca^2+^, PIP_2_ forms stable electrostatic interactions within 5 Å of basic residues on α6. Specifically, the phosphate at position 5 (P5) of the PIP_2_ inositol ring interacts with R665, while the P1-phosphate interacts with K661 (**Fig. 1fi–h**). When Ca^2+^ binds, the α4 helix reorients, causing an immediate disengagement of P5 and R665 and a more gradual disengagement of P1 and K661, which is fully disrupted after approximately 700 ns into the GMD simulation (**Fig. 1g–h**). In place of these interactions, the P4-phosphate of PIP_2_ immediately engages R562 α4 and subsequently K566 after ~800 ns (**Fig. 1fii– j**).

To validate this dynamic model, we performed patch-clamp and measured currents from TMEM16A channels with point mutations in the putative PIP_2_ interaction interface. Wild-type mouse TMEM16A channels expressed in HEK293T cells produced mean currents of 145 ± 26 pA/pF (mean ± SD, n = 60) at 100 mV with 455 nM free Ca^2+^ in the intracellular recording buffer (**Fig. 2a**) ^12^. Mutating R562 to glutamine (R562Q) had no effect on current magnitude, while substituting R562 with alanine (R562A) abolished channel activity. Interestingly, substitutions at K566 enhanced channel activity. Substituting K566 with alanine (K566A) or glutamine (K566Q) resulted in 3.2- and 3.8-fold increases in mean current, respectively (**Fig. 2a**).

**Figure 2:**
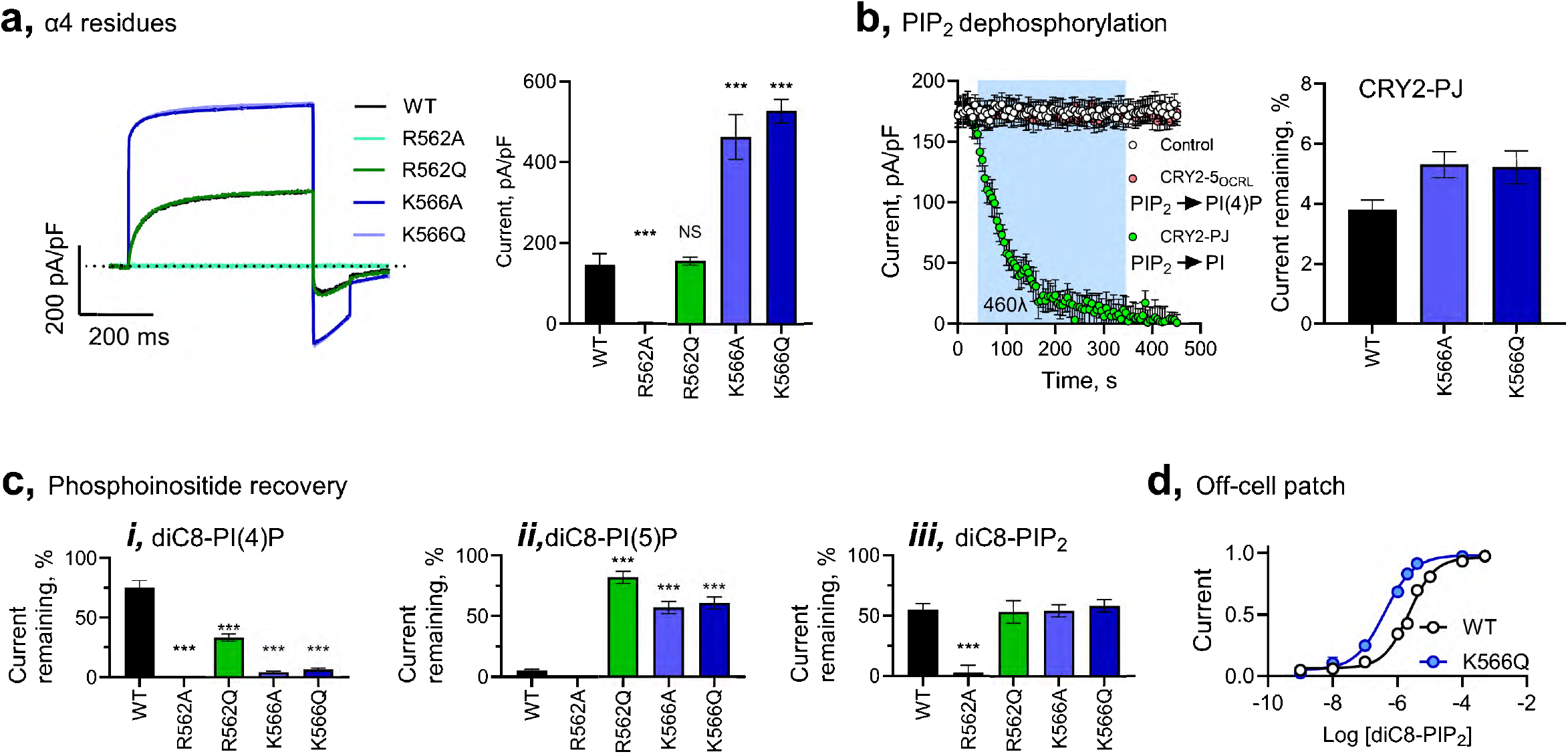
The functional significance of interactions with PIP_2_ in the orthosteric site. mTMEM16A was expressed in HEK293T cells and studied in whole-cell or excised inside-out patch recordings. The electrophysiological data are presented as mean ± standard deviation (S.D.) for at least 8 independent recordings per condition. Statistical differences between the groups was determined by two-tailed, unpaired, Students t-test. *** denotes P<0.001; NS is not significant. **a**, *Left*: Example whole-cell TMEM16A currents evoked by a 500 ms step to 100 mV. *Right*: Summary mean current densities from TMEM16A variants. **b**, *Left*: The time course of whole-cell TMEM16A current magnitude following photostimulation (blue box) of the light activated phosphatases CRY2-5OCRL or CRY2-PJ. *Right*: Summary plots of mean current remaining following activation of CRY2-PJ. **c**, Summary of current remaining following photostimulation of TMEM16A with (***i***) diC8-PI(4)P; (***ii***) diC8-PI(5)P, or (***iii***) diC8-PIP_2_ in the recording pipette. **d**, Application of diC8-PIP_2_ to the inside of PIP_2_-depleted off cells patches shows concentration-dependent current with an EC50 of 2.3 µM for wild type TMEM16A and 0.3 µM for TMEM16A-K566Q channels. *Related to Supplemental Figure S2.*

Next, we used light-activated phosphoinositide (PI) phosphatases to investigate the interaction between TMEM16A and specific phosphates on the PIP_2_ headgroup. This robust optogenetic platform is based on blue light (~440–490 nm) photoactivation of Cryptochrome 2 (CRY2) and its protein partner, CIBN ^18^. By fusing CIBN to the membrane anchor CAAX, CRY2 and its cargo are recruited to the inner leaflet of the cell membrane upon photostimulation ^19,20^. Previous studies, employing real-time measurements of PIP_2_-regulated channel activity or PIP_2_ membrane biosensors such as the PH domain of phospholipase-δ, have demonstrated that CRY2-fused PI-phosphatases localize to the membrane within seconds of photoactivation, triggering dephosphorylation of PIP_2_ within seconds. Although PIP_2_ can be resynthesized within minutes, we mitigate its regeneration with continuous photostimulation ^21–24^.

First, we assessed the impact of photoactivation of CRY2 fused to pseudojanin (CRY2-PJ) on TMEM16A currents. Pseudojanin is a fusion construct combining the inositol polyphosphate 5-phosphatase E and the 4-phosphatase Sac1, which depletes PIP_2_ by sequentially generating PI(4)P and PI ^25^. Photoactivation of CRY2-PJ reduced currents from wild-type TMEM16A, as well as K566A and K566Q mutant channels, by approximately 95% over three minutes (**Fig. 2b**). In contrast, photoactivation of the P5-specific phosphatase CRY2-5OCRL, which generates PI(4)P, did not diminish wild-type TMEM16A currents. This result suggests that either PIP_2_ or PI(4)P are required to gate TMEM16A channels and is consistent with our *in silico* model of TMEM16A gating (**Fig. 1b**).

Next, we photoactivated CRY2-PJ and measured the decrease in currents in cells with soluble, short-chain (diC8) PI(4)P, PI(5)P, or PIP_2_ included in the recording pipette. In wild-type channels, diC8-PI(4)P preserved 75 ± 6% (mean ± SD, n = 10) of the current. In contrast, diC8-PIP_2_ preserved 55 ± 5%, while diC8-PI(5)P failed to preserve any current consistent with the predictions of the model that P4 interactions underpin PIP_2_-induced activation in the wild-type channel (**Fig. 2c**). TMEM16A-R562A channels showed no rescue by any of the diC8 PIPs, underscoring the importance of this residue in PIP_2_-induced activation. In contrast, for TMEM16A-R562Q channels, diC8-PI(4)P did rescue 44 ± 3% of the current, diC8-PI(5)P preserved 82 ± 5%, and diC8-PIP_2_ preserved 54 ± 9%, suggesting that in the R562Q mutant the P5 interactions dominate activation of the channel. Interestingly, diC8-PI(4)P did not rescue currents in TMEM16A-K566A or K566Q channels. In contrast, diC8-PI(5)P rescued approximately 60% of the current in the K566 variants, while diC8-PIP_2_ rescued approximately 50% (**Fig. 2c**). Thus, in the K566A/Q channels the P5 interactions dominate activation of the channel.

We then compared the effects of photoactivation of CRY2-5OCRL alone and together with CRY2-4Sac1 and we could demonstrate sensitivity to dephosphorylation of P5 of PIP_2_ (**Fig. S2a**), consistent with the rescue experiments by PI(5)P (**Fig. 2c**). Next, we tested whether P5 or P1 interacting residues in the α6 helix (R665Q and K668Q, respectively) could be rescued by specific PIPs in the patch pipette (**Fig. S2c-e**). Again, PI(5)P and (or PIP_2_) but not PI(4)P could rescue approximately 50% of the two α6 Gln residue mutants (**Fig. S2c-e**).

To investigate the mechanism by which the PIP_2_ phosphates enhanced current through TMEM16A-K566Q channels, we generated a molecular model of TMEM16A-K566Q in the presence of PIP_2_ and bound Ca^2+^ (**Fig. S2f**). In this model, the glutamine substitution at position 566 disrupts the electrostatic interaction with P4, reorienting PIP_2_ within its binding pocket. Additionally, unlike the wild-type conformation, TMEM16A-K566Q exhibits stable contacts between P4 through R562 on α4, as well as P5 through R665 on α6. Moreover, the P1-phosphate establishes a more stable interaction with K668 on α6 compared to the wild-type channel. Overall, in this mutant channel, PIP_2_ forms stable interactions with both α4 and α6 residues, consistent with the observed increase in TMEM16A-K566Q channel activity (**Fig. 1g-i** and **Fig. S2f–h**).

To determine whether the increased current in TMEM16A-K566Q channels could be attributed to an enhanced apparent affinity for PIP_2_, we recorded currents in excised inside-out patches. First, endogenous PIP_2_ was chelated using poly-L-lysine, abolishing channel activity. Subsequently, exogenous diC8-PIP_2_ was applied to the cytosolic face of the membrane. In wild-type TMEM16A channels, diC8-PIP_2_ activated the channels with a half-maximal effective concentration (EC50) of 2.3 µM, consistent with prior studies ^17^. In contrast, diC8-PIP_2_ activated TMEM16A-K566Q channels with an EC50 of 0.3 µM, indicating a tenfold higher apparent affinity for PIP_2_ in the mutant channels (**Fig. 2d**).

### PIP_2_ acyl chains interact with α4

In addition to studying interactions between the phosphatidylinositol headgroup of PIP_2_ and TMEM16A, we examined the interactions between the acyl chains of PIP_2_ and the channel in the Ca^2+^-free and Ca^2+^-bound versions of the site-1 GMD system. Carbons in the acyl chains of PIP_2_ interact with residues on the α4-helix of TMEM16A. These interactions are typically more stable in the Ca^2+^-bound system (**Fig. 3a–b**). To assess the functional significance of this interaction network, we performed patch-clamp experiments on mutants of α4 residues from A542 to L553 designed to reduce the hydrophobic character of each residue. Mutations in residues A542 to I545, located above the acyl chain interaction region, did not affect the current magnitude. In contrast, TMEM16A-N546A and L547A mutants exhibited no measurable current (**Fig. 3c**), which could not be rescued by applying diC8-PIP_2_ or long-chain (LC)-PIP_2_ to the cytosolic face of the membrane in excised patches (**Fig. 3d**). TMEM16A-V548A, V549A, and I550A mutants exhibited approximately 50% of the wild-type current and these currents were not enhanced by diC8-PIP_2_. For TMEM16A-V548A, no additional recovery was observed when the channels were studied in excised patches and exposed to LC-PIP_2_. In contrast, TMEM16A-I550A channels whose activity was approximately half of the wild-type showed robust recovery, achieving current levels like wild-type in response to LC-PIP_2_ but did not respond further to diC8-PIP_2_. Concentration-response experiments revealed that the reduced current in TMEM16A-I550A channels could be attributed to a lower apparent affinity for diC8-PIP_2_, with an EC50 of 11 µM— approximately 5-fold lower than the wild-type EC50 (**Fig. 3d**). I551A showed an approximately 75% remaining current relative to control, which could be fully recovered by either diC8- or LC-PIP_2_. Alanine substitutions for residues L552 and L553 produced channels with similar behavior to the wild type (**Fig. 3b-d**). These findings define the N546 to I551 region of α4 as a critical determinant of acyl chain action, supporting our *in silico* model and demonstrating the importance of the PIP_2_ acyl chains in mediating TMEM16A gating.

**Figure 3:**
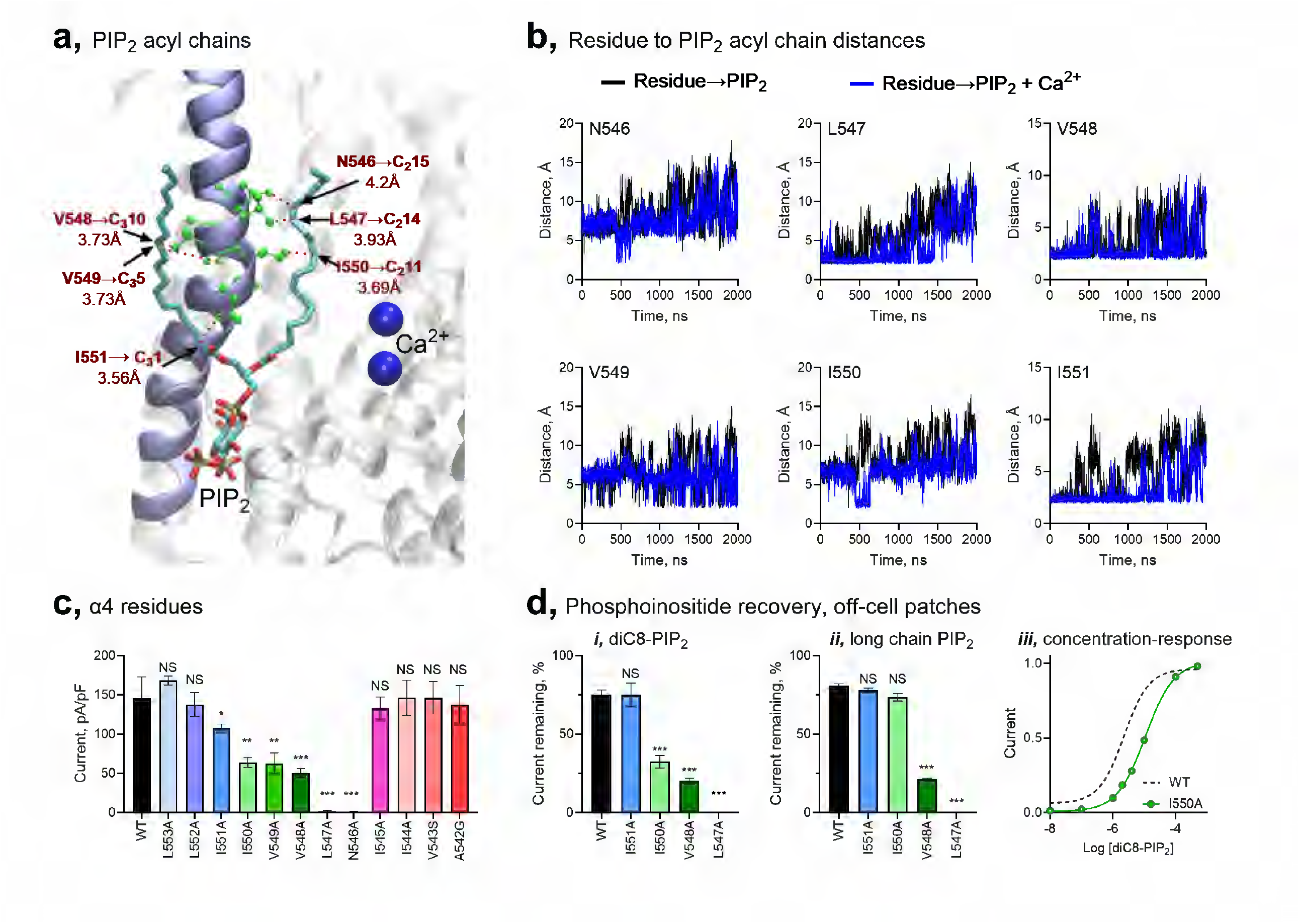
The PIP_2_ acyl chain interact with α4. Site-1 GMD studies show the PIP_2_ acyl chain carbon interactions with α4 residues N546 through to I551. The electrophysiological data are presented as mean ± standard deviation (S.D.) for at least 8 independent recordings per condition. Statistical differences between the groups were determined by two-tailed, unpaired, Students t-test. * denotes P<0.05, **P<0.01 and *** P<0.001; NS is not significant. **a**, Interaction distances between the residues and the closest PIP_2_ carbon are noted. **b**, Trajectory plots show that interactions are stabilized in the presence of PIP_2_ and Ca^2+^. **c**, Summary of current magnitudes when residues in the α4 are substituted for alanine. **d**, Summary of current rescue in off-cell patches by the application of (***i***) diC8-PIP_2_ or (***ii***) long-chain PIP_2_; (***iii***) The apparent affinity of TMEM16A-I550A to diC8-PIP_2_ is 11 µM, compared to 2.3 µM for the wild-type channel.

### An electrostatic gate in the corpus of TMEM16A

The exact location and structure of a limiting constriction, or gate, within the TMEM16A permeation pathway remains unclear. Our GMD simulations of the Ca^2+^-bound site-1 system reveal that a narrow region of the ion permeation pathway is formed by a ring of residues—D554 (α4), K588 (α5), K645 (α6), and E705 (α7)—located at the center of the channel. D554 is located in the α4 helix that PIP_2_ is acting on to activate the channel. Its strategic position between PIP_2_-interacting residues, I551 on one side with the acyl chains of PIP_2_ and R562 on its other side with P4 of the PIP_2_ headgroup. E705 is part of the channel’s Ca^2+^-binding pocket. In the presence of PIP_2_ but the absence of Ca^2+^, the aperture formed by these residues is too narrow to permit Cl^−^ permeation through the channel. Upon Ca^2+^ binding, E705 shifts to participate in the coordination of Ca^2+^, inducing a rotation of K645 that increases the diameter of the aperture, allowing Cl^−^ ions to pass through the channel (**Fig. 4a**). These observations suggest that the electrostatic ring serves as a gating mechanism within the TMEM16A channel.

**Figure 4.**
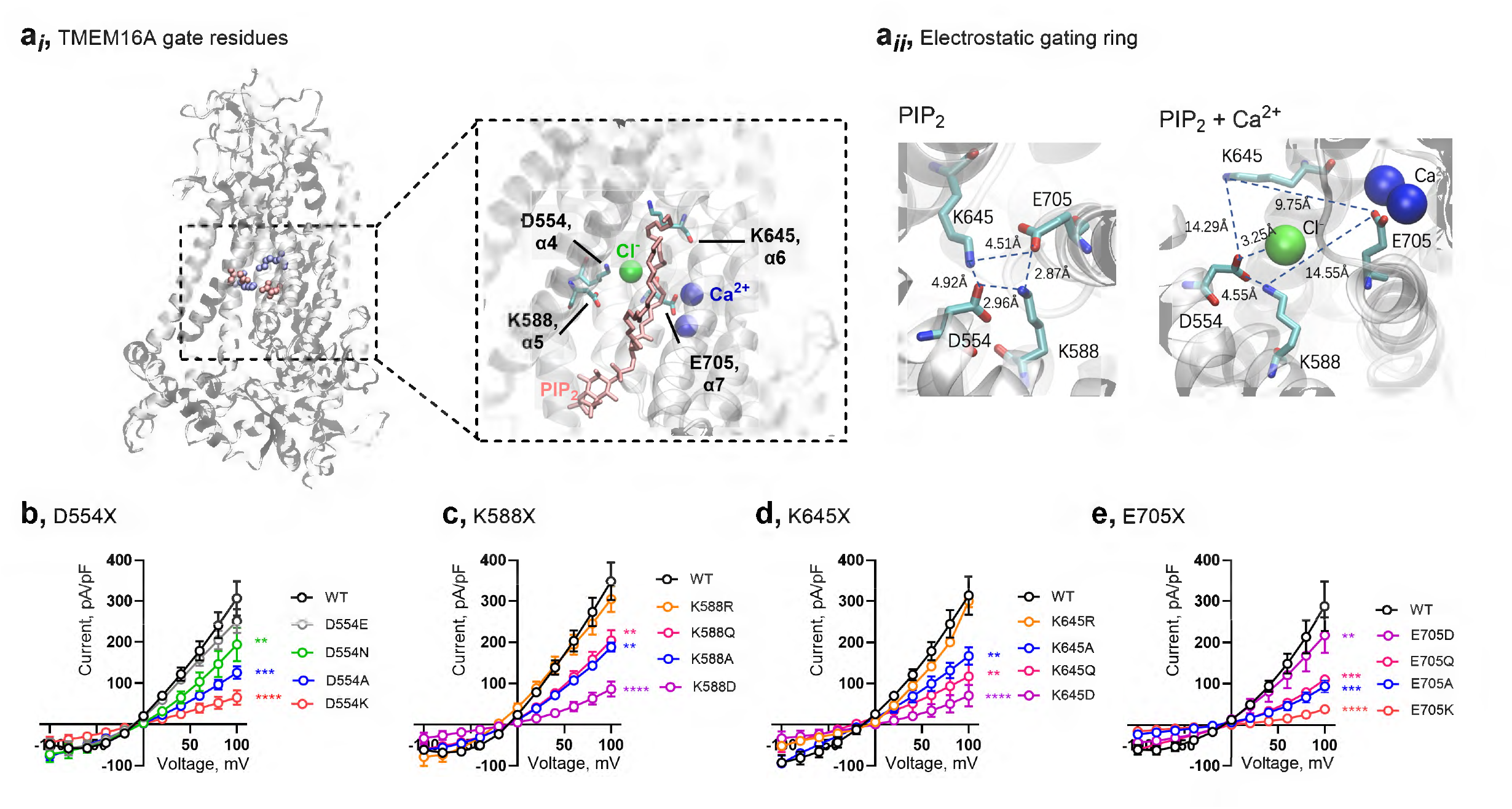
An electrostatic gating ring in the corpus of TMEM16A. The electrophysiological data are presented as mean ± standard deviation (S.D.) for at least 8 independent recordings per condition. Statistical differences between the groups were determined by two-tailed, unpaired, Students t-test. ** denotes P<0.01, ***P<0.001, **** P<0.0001. **ai**, Snapshot from the site-1 GMD study showing the relative position of D554 (α4), K588 (α5), K656 (α6) and, E705 (α7) in the presence of PIP_2_ and Ca^2+^. **aii**, Snapshots of the gating ring showing the distances between the residues in the presence of PIP_2_ (*left*) and PIP_2_ with Ca^2+^, where Cl-is shown permeating the channel (*right*). **b**, Mean current-voltage plots for wild type TMEM16A and D554 variants. **c**, Mean current-voltage plots for wild type TMEM16A and K588 variants. **d**, Mean current-voltage plots for wild type TMEM16A and K645 variants. **e**, Mean current-voltage plots for wild type TMEM16A and E705 variants.

To validate this prediction, mutagenesis studies were designed to determine the role of the electrostatic ring as a gate. Conserving the native charge of these residues had small (in E705D reduced current magnitude by approximately 25% at 100 mV (n = 11, P < 0.01)) to no effects on current magnitude (in D554E, K588R, and K645R channels). In all cases, reversing the charge of the residues reduced the current magnitude by over 90%. Substitutions to alanine or neutral residues with conserved volume reduced current magnitude by ~50% in K588Q/A, K645Q/A, and E705Q/A channels. Similarly, D554N exhibited a ~50% reduction in current magnitude, whereas D554A displayed a more pronounced reduction of ~60% compared to the wild-type channel (**Fig. 4b–e**).

### The extracellular mouth of the TMEM16A pore

Last, we investigated the effect of PIP_2_ on the α4 extracellular domain of the channel by examining the trajectory of interactions between the flexible α4 and the α6 helices in the Ca^2+^-bound site-1 system. PCA revealed that the external portion of α4 undergoes significant fluctuations in motion upon Ca^2+^ binding (**Fig. 1c–e**). In the absence of PIP_2_, the α4-residue R535 can form a salt bridge with E633 on α6. In the presence of PIP_2_ alone these two residues stably move away from each other precluding their interaction. Moreover, when both PIP_2_ and Ca^2+^ are present, the α4 helix pivots away from α6 by approximately 20 Å (**Fig. 5a, b**). This conformational change, driven by PIP_2_ interactions with α4 depletes the accumulated cloud of Cl^−^ ions at the extracellular mouth of TMEM16A as Cl^−^ ions enter the permeation pathway of the channel (**Fig. 5c**). Supporting these modeling predictions, whole-cell patch-clamp recordings show that mutations in any of three residues, where Cl^-^ was predicted to accumulate, significantly enhance Cl^−^ currents through TMEM16A channels (**Fig. 5d, e**). **Figure 5f** compares the human and mouse amino acid sequences of the α4 helix and summarizes the residue interactions of the PIP_2_ headgroup phosphates and acyl chain carbons on either side of the D554 residue of the channel’s gating ring. Residues in the extracellular side of the α4 helix marking the Cl^-^ entry to the permeation pathway are also shown.

**Figure 5.**
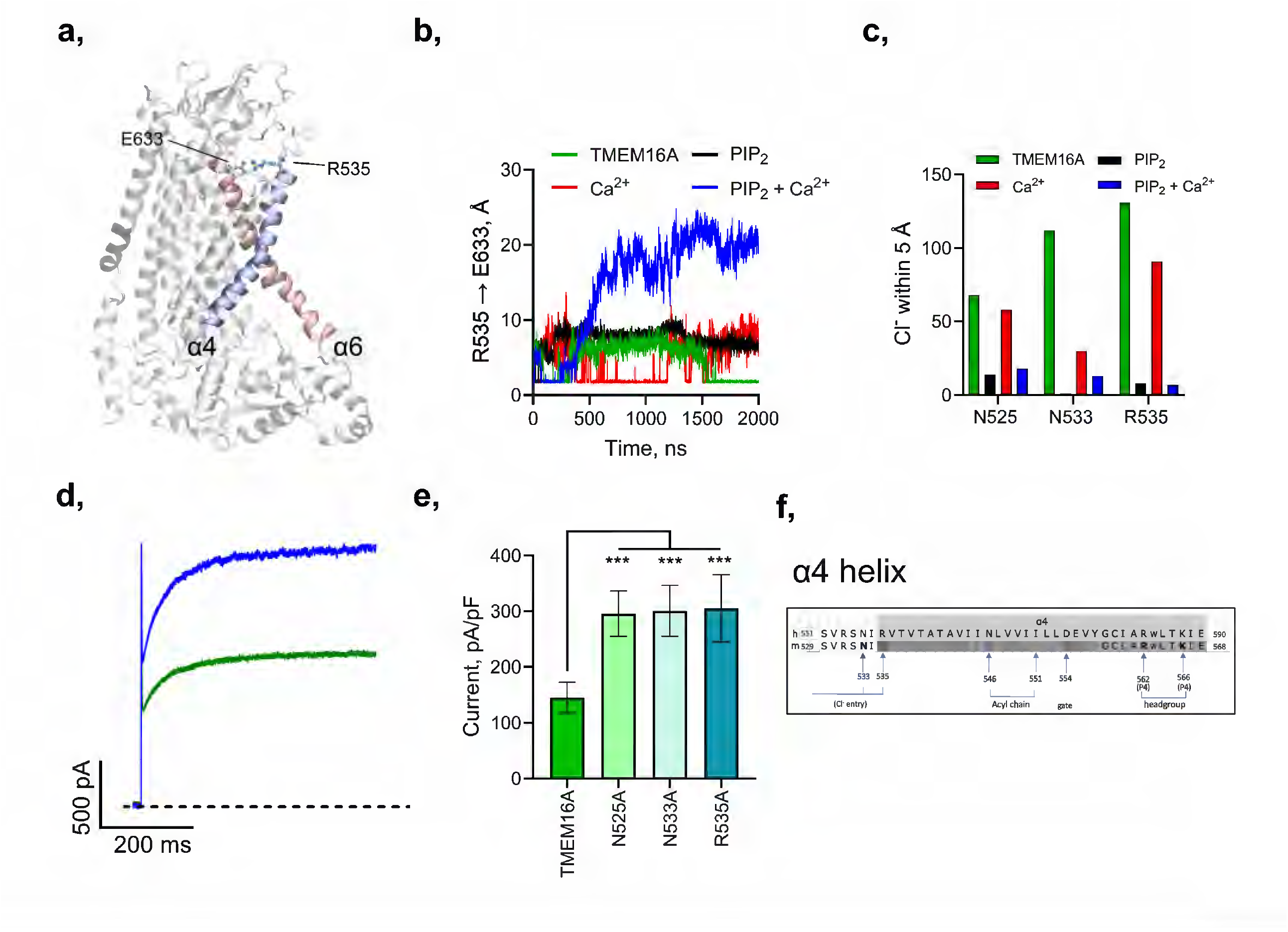
PIP_2_ is required to open the external mouth of the pore. The electrophysiological data are presented as mean ± standard deviation (S.D.) for at least 8 independent recordings per condition. Statistical differences between the groups was determined by two-tailed, unpaired, Students t-test. *** denotes P<0.001. **aa**, Model showing a salt bridge between R535 (α4) and E633 (α6). **b**, Trajectory plot showing disruption in the R535 to E633 interaction with PIP_2_ and Ca^2+^ bound to the channel. **c**, Summary of the number of Cl-ions retained within 5Å of the residue noted over 2000 ns simulations. **d**, Example whole-cell currents for TMEM16A wild type (green) and TMEM16A-R535 channels. **e**, Summary of mean current-densities for the wild-type channel and the variants shown. **f**, Schema showing the location of the major residues on the α4 helix that interact with PIP_2_ or form part of the gating ring structure (i.e., D554) or the entry point of the permeation pathway (e.g., R535).

## Discussion

In this study, we identify and characterize dynamic PIP_2_ binding interactions required to gate the TMEM16A ion channel. By analyzing GMD simulation trajectories in a docking-model that allows Cl^-^ permeation, we found critical contacts between TMEM16A and the headgroup phosphates and acyl chain carbons of PIP_2_. Targeted electrophysiology experiments validated our model, confirming the location of a dynamic orthosteric PIP_2_ binding pocket between α4 and α6 that controls the TMEM16A gating machinery.

Our study elucidates the Cl^−^ permeation pathway through individual TMEM16A subunits and provides critical insights into the role of PIP_2_ in facilitating channel activation. In the basal state, the P1 and P5-phosphates on the PIP_2_ headgroup interact with specific residues on the α6 helix. Upon Ca^2+^ binding, the pocket undergoes a conformational reorientation, initiating interactions between the P4-phosphate and the α4 helix. This structural rearrangement leads to Cl^-^ permeation by dilating a ring of charged residues that form an electrostatic gate deep within the core of each subunit and promoting the displacement of the N-terminal region of the α4 helix, resulting in a widened extracellular pore entrance that enhances Cl^−^ flux.

In addition to elucidating the role of each phosphate in the PIP_2_ headgroup, we identify specific interactions between the PIP_2_ acyl chain carbons and the α4 helix that participate in channel activation. Thus, as Ca^2+^ binds TMEM16A and the PIP_2_-headgroup binding pocket interactions reorient, interaction stability between a stretch of hydrophobic residues on α4 and the acyl chain carbons of PIP_2_ increases. Alanine substitutions between N546 and I551 impair channel function. It is noteworthy that the reduced current phenotype of TMEM16A-I550A is partially rescued by short-chain diC8-PIP_2_, and completely by long-chain PIP_2_, which can form additional interactions with α4, overall. D554 located on α4, just below the hydrophobic stretch of residues. is one of the four residues that forms the electrostatic gating ring in the corpus of the channel that constricts the Cl^-^ permeation pathway in the absence of Ca^2+^. When Ca^2+^ binds, the structural rearrangement of the channel enlarges the distance between D554 on α4, and the opposite gating ring residue, E705 on α7 by approximately 2.6-fold and the distance between K588 (α5) and K645 (α6) by approximately 2.3-fold, enlarging the aperture and allowing Cl^-^ ions to pass. This is, to the best of our knowledge, the first example of an electrostatic gating ring mechanism in the interior of an ion channel.

Previous studies have proposed numerous PIP_2_ binding sites on TMEM16A, many of which show partial overlap with the putative binding interfaces we observed in our site-2, 3, and 4 models ^10,11,16,17^. These alternative sites do not support Cl^−^ flux *in silico*. Mutagenesis studies targeting these sites, as reported previously, have demonstrated changes to channel activity, though such effects are likely allosteric in nature. In contrast, our findings underscore that the interactions between PIP_2_ and site-1 are both necessary and sufficient for TMEM16A gating. This conclusion is substantiated by multiple lines of evidence. Our GMD models for site-1 exhibited a strong concordance with our experimental data, including the restoration of TMEM16A activity via exogenous application of diC8 or long-chain PIP_2_ in membranes depleted of endogenous PIP_2_. Furthermore, our model predicted that specific mutations would evoke changes in the activity of TMEM16A that were then captured in electrophysiological experiments.

PIP_2_ is the most abundant phosphoinositide in the plasma membrane and is continuously synthesized through the phosphorylation of PI(4)P by PI(4)P-5-kinases^26^. Although PI(4)P is a comparatively minor species, our studies reveal that it can support TMEM16A activity. This functionality was demonstrated through the production of PI(4)P via CRY2-5POCRL-mediated dephosphorylation of PIP_2_ and the exogenous application of diC8-PI(4)P to PIP_2_-depleted cells. Notably, we found that the phosphate group at position 5 (P5) of PIP_2_ does not play a significant role in TMEM16A gating upon Ca^2+^ binding. In contrast, the phosphate group at position 4 (P4) interacts robustly with residues R562 and K566 on the α4 helix. Physiologically, depletion of PIP_2_ typically results from phospholipase C hydrolysis, downstream of Gαq signaling. This signaling cascade generates IP3-induced increases in cytosolic Ca^2+^ levels which activate TMEM16A. However, these currents deactivate rapidly as PIP_2_ levels in the membrane decrease. Given that PI(4)P is both a substrate for PIP_2_ and can also support TMEM16A currents, the relative rates and the extend of TMEM16A current activation and deactivation in response to Gαq signaling will be contingent upon the relative abundance of PI(4)P, which varies across cell types ^25,27,28^.

During the GMD simulations only one TMEM16A subunit conducted Cl^−^ ions. This facet of the model requires further study to determine if the two Cl^-^ conductance pathways in the channel operate in a mutually exclusive manner. Of note, classical CLC chloride channels form as dimers of subunits that function independently in a “double-barreled shotgun” configuration ^29,30^. Our computational model paves the way for future investigations into the dynamic inter-subunit interactions governing TMEM16A function.

Overall, this study demonstrates that GMD simulations, coupled with functional validation, serve as an effective strategy to dissect molecular mechanisms that underlay PIP_2_-enabled ion channel gating. By confirming the predictions of our *in-silico* study, we show that our GMD model can provide a robust platform to unravel additional questions about the structural dynamics that govern the activity of TMEM16A channels. This approach has potential implication for dissecting the mechanisms underlying modulators of the channels and predicting structure-activity relationships for drug development.

## Supplemental Figures

**Figure S1: Nonfunctional PIP2 binding sites on TMEM16A.** PIP_2_ was docked to the Ca^2+^-bound TMEM16A channel (PDB: 5OYB) and studied for *in silico* conduction of Cl^-^ by gating molecular dynamics. The Site-2, -3 and -4 systems bound PIP_2_ but did not yield Cl^-^ permeation.

***Related to Figure 1***.

**Figure S2: TMEM16A-K566 variant channels**. The data are means ± standard deviation (S.D.) for at least 8 independent recordings per condition. Statistical differences between the groups was determined by two-tailed, unpaired, Students t-test. *** denotes P<0.001. **a**, mTMEM16A-K566Q/A channels were expressed in HEK293T cells and the time course of changes in whole-cell current magnitude was measured following photostimulation (blue box) of the light activated 5-phosphatase CRY2-5OCRL or CRY2-5OCRL + the 4-phosphatase, CRY2-Sac1 which were co-expressed in the cells. **b**, Summary plots of mean current remaining following photostimulation of phosphatases for the type, R665Q and K668A channels. **c**, Summary of current remaining following photostimulation of TMEM16A with diC8-PI(4)P; **d**, diC8-PI(5)P, or **e**, diC8-PIP_2_ in the recording pipette. **f**, Binding pose of PIP_2_ in site-1 of the TMEM16A-K566Q channel in the presence of Ca^2+^. Comparison of distances of P1, P4, and P5 phosphates from α6 and α4 residues for TMEM16A WT versus K566Q channels: **g*i***, Trajectories of interaction between P5 and R665; ***ii***, P1-K661; ***iii***, P1-K668; ***iv***, P4-R562 and ***v***, P4-K/Q566. **h**, Summary of mean current-densities for TMEM16A-α6 variants shown.

***Related to Figures 1 and 2***.

## Methods

### Molecular modelling and docking

The cryo-EM structure of mouse TMEM16A (PDBID: 5OYB) was prepared by the Protein Preparation Wizard module of Maestro (2019-3) program (Schrödinger, Inc. New York, NY) and subsequently utilized for docking. To systemically search the potential PIP_2_ binding sites, we covered the channel with 56 grid boxes (each with centers spaced 20 Å apart) using an in-house script. Induced-fit-docking (IFD) of the Glide program from Schrödinger (Schrödinger, Inc. New York, NY) was used for docking simulations ^31^. The diC1 PIP_2_ was initially put at the center of the boxes to conduct each docking simulation, using the standard precision (SP) module in the Glide program ^32^. The default parameters were used for IFD simulations. The residues within 3.5Å of ligand poses were selected for side chain optimization by prime refinement. The SP score was used for ranking the ligand poses. The pose of docked ligand with the lowest docking SP score was selected for each potential binding site. Four predicted binding sites with the lowest docking SP scores were selected, and the diC1 PIP_2_ was replaced with a full-length PIP_2_ for the MD simulations.

### System set-up and molecular dynamics simulations

Protonation states of the titratable residues in TMEM16A channel were calculated at pH = 7.4 via the use of the H++ server (http://biophysics.cs.vt.edu/) ^33^. The simulation system was generated using the CHARMM-GUI Membrane Builder webserver (http://www.charmmgui.org/?doc=input/membrane) ^34–36^. First, the protein was inserted in an explicit lipid bilayer of POPC, POPE, POPS and cholesterol with molecular ratio of 25:5:5:1 8 ^37^. Then, the complexes were put in a water box (176Å×176Å×156Å), followed by an addition of 150 mM KCl to the system.

Molecular dynamics simulations were conducted using the PMEMD.CUDA program in AMBER169. The tleap module was employed to neutralize the system by extra K^+^ or Cl^−^ ions as needed. The FF14SB, LIPID17, and GAFF force fields were chosen for protein, mixed lipid membrane and PIP_2_, respectively. The parameters of PIP_2_ were generated using the general AMBER force field by the Antechamber module of AmberTools 17 and using the partial charge determined via restrained electrostatic potential charge-fitting scheme by ab initio quantum chemistry at the HF/6-31G* level. The systems were energetically minimized with 2,000 steps of steepest descent followed by 3,000 steps of conjugate gradient. Subsequently, Langevin dynamics was used to heat the systems from an initial temperature of 0 K to a final temperature of 303 K, with a collision frequency of 1 ps^−1^. During heating, an initial constant force of 500 kcal·mol^−1^·Å^−2^ was applied to positionally restrain the receptor complexes, which was then gradually reduced to 10 kcal·mol^−1^·Å^−2^, allowing the lipid and water molecules to move freely. Subsequently, the systems underwent 5 ns of equilibrium molecular dynamics simulations. Finally, a total of 2 μs of production molecular dynamics simulations were conducted, with coordinates saved every 100 ps for subsequent analysis. The simulations were conducted with periodic boundary conditions, initially in a constant temperature, constant-pressure ensemble (NPT), followed by a constant-temperature, constant volume ensemble (NVT). The pressure was regulated using the isotropic position scaling algorithm, with the pressure relaxation time set to 2.0 ps. The particle mesh Ewald (PME) method was used to calculate the long-range electrostatics with a 10 Å cut-off ^38^. An external voltage of 0.06 V/nm (approximately 200mV across the membrane) was added to the systems from the extracellular to the intracellular side^24,39,40^. A 4-fs time step was employed using the hydrogen mass repartition algorithm for systems to accelerate the molecular dynamics simulations ^41^.

### Analysis of MD runs

Distance analysis and principal component analysis (PCA) were performed using GROMACS 5.1.4 ^42^. PCA was performed to extract the collective motions of the channel from the MD simulation trajectories. This describes the motions with a set of eigenvector and eigenvalue pairs, which are obtained by diagonalizing the covariance matrix of the Cα atomic positional fluctuations^43,44^. The analysis program gmx_anaeig within GROMACS was used to conduct PCA. The interaction between TMEM16A and PIP_2_ was analyzed by Discovery Studio 2017 software. VMD 1.9.3 was used for structure rendering ^45^.

### Molecular biology and reagents

Purified diC8-PIP_2_, PI(4)P and PI(5)P and long-chain (LC) PIP_2_ were purchased from Echelon Biosciences. CIBN-CAAX and mCherry-CRY2-5POCRL were gifts from the De Camilli lab (Yale University, New Haven, CT). The open reading frame of pseudojanin ^25^ was designed with an N-terminal CRY2-tag in the pMAX(+) vector and was generated by Genscript (Piscataway, NY). Mouse TMEM16A tagged with eGFP in the peGFP vector was a gift from Lily Jan (UCSF). Mutations were introduced by Quikchange mutagenesis according to the manufacturer’s protocols.

### Cell culture and transfection

Human embryonic kidney (HEK293T) cells were acquired from American Type Culture Collection (ATCC, Cat #CRL-3216) and were maintained in Dulbecco’s modified Eagle’s medium (ATCC) supplemented with 100 units/ml penicillin, 100 μg/ml streptomycin, and 10% (vol/vol) fetal bovine serum. Cells were routinely confirmed to be mycoplasma free via PCR and Hoescht stain. The cells were incubated in a 37°C, humified incubator supplemented with 5% CO2. For our experiments, cells were seeded on glass coverslips in 35 mm culture dishes at least one day before transfection. Cells were transiently transfected with TMEM16A variants, CIBN-CAAX and CRY2-5POCRL or CRY2-pseudojanin as indicated, in OptiMEM using polyethyleneimine (PEI) for 1-2 hours at a ratio of 1 μg of DNA to 4 μL PEI. The near red fluorescent protein, mCherry, was typically used as a transfection marker. The cells were studied 24-48 hours after transfection.

### Patch clamp recording

HEK293T cells were studied by whole cell patch-clamp. Currents were recorded with a Tecella Pico-2 amplifier (Tecella) controlled using WinWCP software (University of Strathclyde). Currents were low pass (Bessel) filtered at 9 kHz and digitized at 50 kHz. Cells were studied in an external recording buffer comprising (in mM): 150 NaCl, 1 CaCl_2_, 1 MgCl_2_, 10 glucose, 10 HEPES, adjusted to pH 7.4 with NaOH. Patch pipettes were pulled from borosilicate glass (Clark Kent), using a vertical puller (Narishige) and had a resistance of 2.5–4 MΩ when filled with internal solution containing (in mM): 130 CsCl, 1 MgCl_2_, 10 EGTA, with 3 Na_2_ATP adjusted to pH 7.35 with CsOH. The recording conditions are based on those described by the Hille and Suh labs and give a reversal potential ECl of −4mV ^12^. In accord with this study, we determined that 8.47mM Ca^2+^ from CaCl_2_ leaves a free Ca^2+^ concentration of 455 nΜ under our experimental conditions, allowing robust activation of TMEM16A channels. The free Ca^2+^ concentration is calculated using the MAXCHELATOR online algorithm housed at UC Davis. For whole-cell studies, the cells are held at −80mV for 2s before stepping to a 500ms test potential between −120mV and 120mV with 10mV increments. If the current-voltage relationship is observed to deviate from a reversal potential of ~−4mV this would indicate a change in the selectivity of the channel. For time course studies, this protocol is modified to step to 100 mV only. Whole-cell currents were normalized to cell capacitance. Mean ± SEM capacitance values were 12 ± 4 pF for HEK293T cells. For off-cell patch experiments, the pipettes were filled with the external recording buffer and the inside face of the patches was perfused with the internal solution and the polarity of the voltage-clamp protocols was inverted. After initial appraisal of the current magnitude, the patches were perfused with poly L-lysine (100 µg/ml) to chelate phosphoinositides from the membrane.

For simultaneous optogenetic-patch clamp studies the blue light system was photoactivated in epifluorescence-mode using continuous excitation from a broad-spectrum LED (Excelitas) through a 448/ 20 nm filter (Chroma), via a 20x objective lens (Olympus). The light output at the sample was measured at 50 mW/cm2 by a photometer (ThorLabs). To avoid pre-activation of CRY2-fusion proteins, cells were incubated in the dark and handled in foil wrapped dishes prior to the experiments. Cells of interest were identified using mCherry as a transfection marker because its spectral properties do not activate CRY2. Further, we use a transilluminator that includes a bandpass filter to block blue light during brightfield visualization. To generate paired data, the currents were recorded first in the dark and again following a 3-min pre-illumination with epifluorescent blue-light that persisted through the remainder of the study. All the experiments were performed at room temperature.

## Statistics

Data were handled in WinWCP, Clampfit, and Excel software with statistical analysis performed using GraphPad (Prism). The data are presented as mean ± standard deviation (S.D.) with statistical differences between paired groups determined by two-tailed, unpaired, Students t-test, unless indicated otherwise. The threshold for significance was determined to be p<0.05.

## Acknowledgements

The work was funded by R01 HL059949-27 to DEL. The authors are grateful to H. Vaananen, AM Baggetta and AK Yauch for technical support and to J Pentikis and H Sui for making most of the mutants used in this study. They are also grateful to IM Vynichaki, RC Kissell and QY Tang for providing functional support for different experiments within the study. The authors thank JL Sui for introducing them to this project and Z Zhang for supporting J Xu to spend a year at Northeastern University when the bulk of the GMD simulations were performed to yield the wild-type model. They also thank C Iliopoulos-Tsoutsouvas, M Gerasi, SM Smirnakis, D Woulfe, and C Miller, for useful comments on the manuscript.

## Author Contributions

J. X., M.Z., N.P., M.C. Performed computational studies and data analysis. A.S.C, A.C., L.D.P. Performed electrophysiological studies and data analysis. T.K. designed and generated plasmid constructs. D.E.L., L.D.P., wrote the paper and prepared the figures. All authors commented on the manuscript.

