## Supplement for "Orthosteric interactions with PIP_2_ activate TMEM16A channels"

**Figure S1: Nonfunctional $PIP_2$ binding sites on TMEM16A.**

**Figure S2: TMEM16A-K566 variant channels.**

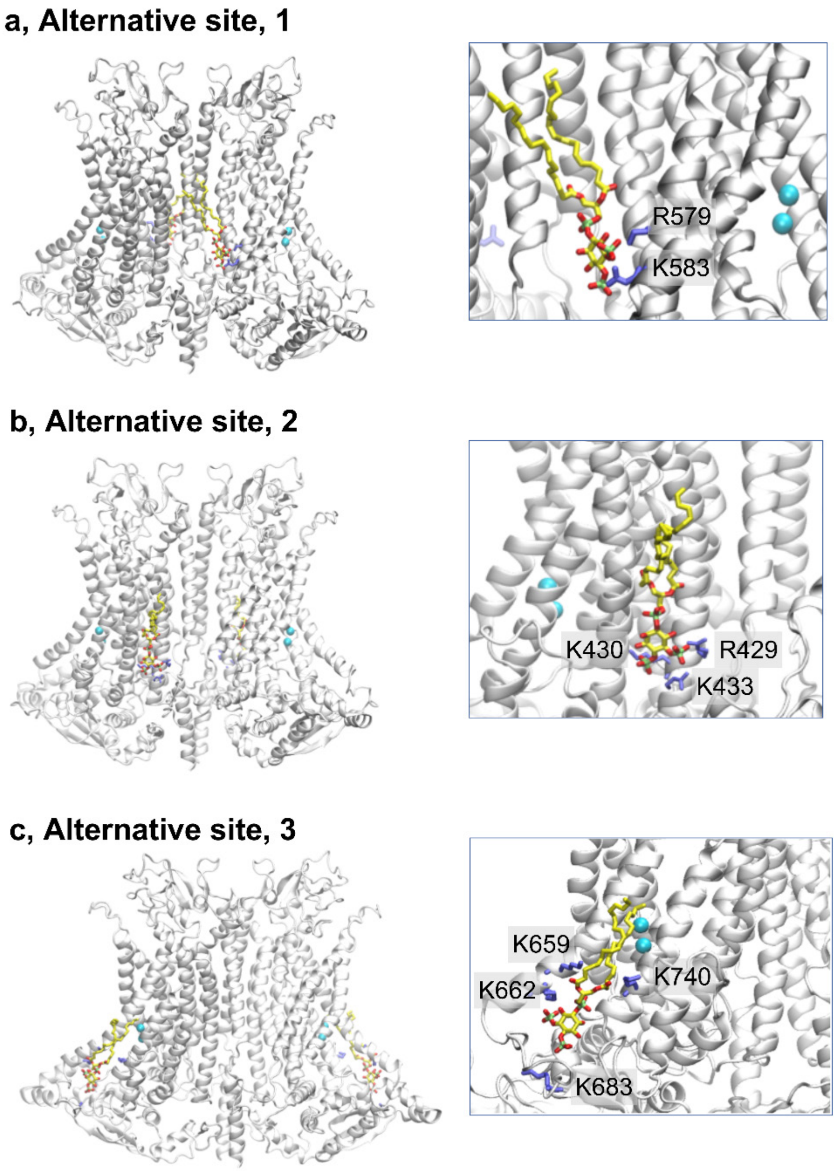


**Figure S1: Nonfunctional $PIP_2$ binding sites on TMEM16A.** $PIP_2$ was docked to the $Ca^{2+}$-bound TMEM16A channel (PDB: 5OYB) and studied for *in silico* conduction of $Cl^-$ by gating molecular dynamics. The Site-2, -3 and -4 systems bound $PIP_2$ but did not yield $Cl^-$ permeation.

***Related to Figure 1.***

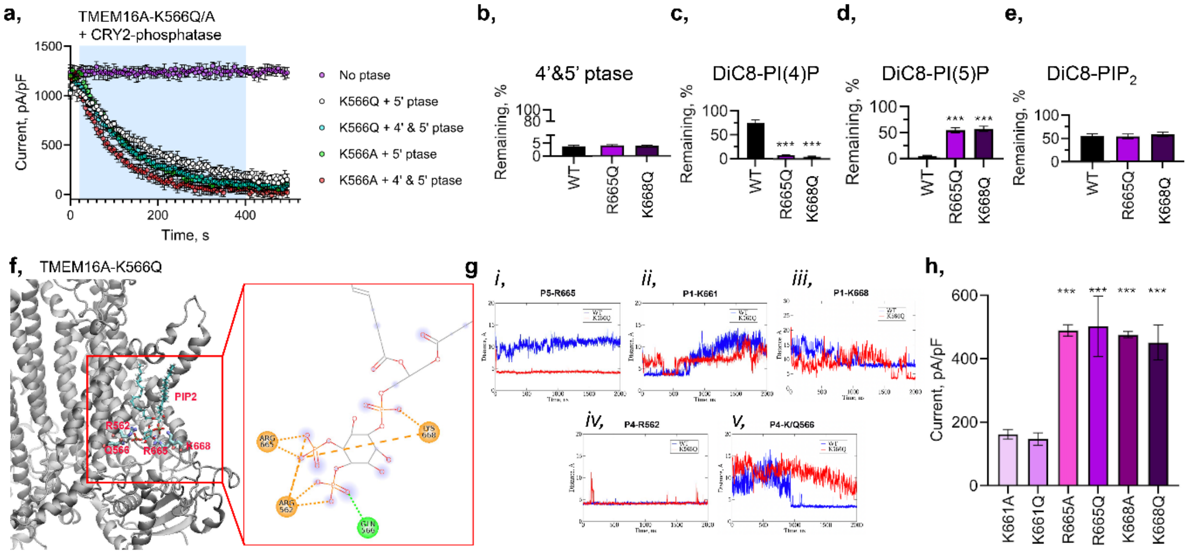


**Figure S2: TMEM16A-K566 variant channels.** The data are means ± standard deviation (S.D.) for at least 8 independent recordings per condition. Statistical differences between the groups was determined by two-tailed, unpaired, Students t-test. *** denotes P<0.001. **a,** mTMEM16A-K566Q/A channels were expressed in HEK293T cells and the time course of changes in whole-cell current magnitude was measured following photostimulation (blue box) of the light activated 5-phosphatase CRY2-5$_{\mathrm{OCRL}}$ or CRY2-5$_{\mathrm{OCRL}}$ + the 4-phosphatase, CRY2-Sac1 which were co-expressed in the cells . **b**, Summary plots of mean current remaining following photostimulation of phosphatases for the type, R665Q and K668A channels. **c**, Summary of current remaining following photostimulation of TMEM16A with diC8-PI(4)P; **d,** diC8-PI(5)P, or **e,** diC8-PIP$_2$ in the recording pipette. **f**, Binding pose of PIP$_2$ in site-1 of the TMEM16A-K566Q channel in the presence of Ca$^{2+}$. Comparison of distances of P1, P4, and P5 phosphates from α6 and α4 residues for TMEM16A WT versus K566Q channels: ***gi***, Trajectories of interaction between P5 and R665; ***ii,*** P1-K661; ***iii,*** P1-K668; ***iv,*** P4-R562 and ***v,*** P4-K/Q566. **h,** Summary of mean current-densities for TMEM16A-α6 variants shown.

***Related to Figures 1 and 2.***
